# A dementia-associated risk variant near *TMEM106B* alters chromatin architecture and gene expression

**DOI:** 10.1101/154989

**Authors:** Michael D. Gallagher, Marijan Posavi, Peng Huang, Travis L. Unger, Yosef Berlyand, Analise L. Gruenewald, Alessandra Chesi, Elisabetta Manduchi, Andrew D. Wells, Struan F.A. Grant, Gerd A. Blobel, Christopher D. Brown, Alice S. Chen-Plotkin

## Abstract

Neurodegenerative diseases pose an extraordinary threat to the world’s aging population, yet no disease-modifying therapies are available. While genome-wide association studies (GWAS) have identified hundreds of novel risk loci for neurodegeneration, the mechanisms by which these loci influence disease risk are largely unknown. Indeed, of the many thousands of SNP-trait associations identified by GWAS over the past ~10 years, very few are understood mechanistically. Here, we investigate the association of common genetic variants at the 7p21 locus with risk for the neurodegenerative disease frontotemporal lobar degeneration. We show that variants associated with disease risk correlate with increased brain expression of the 7p21 gene *TMEM106B*, and no other genes. Furthermore, incremental increases in TMEM106B levels result in incremental increases in lysosomal phenotypes and cell toxicity. We then combine fine-mapping, bioinformatics, and bench-based approaches to functionally characterize all candidate causal variants at this locus. This approach identified a noncoding variant, rs1990620, which differentially recruits CTCF, influencing CTCF-mediated long-range chromatin looping interactions between multiple *cis*-regulatory elements, including the *TMEM106B* promoter. Our findings thus provide an in-depth analysis of the 7p21 locus linked by GWAS to frontotemporal lobar degeneration, nominating a causal variant and a causal mechanism for allele-specific expression and disease association at this locus. Finally, we show that genetic variants associated with risk for neurodegenerative diseases beyond frontotemporal lobar degeneration are enriched in brain CTCF-binding sites genome-wide, implicating CTCF-mediated gene regulation in risk for neurodegeneration more generally.

## INTRODUCTION

Neurodegenerative diseases are a leading cause of disability and death in the developed world, with numbers affected by these diseases poised to increase as the world population ages. There are still no disease-modifying therapies for the major late-onset neurodegenerative diseases such as Alzheimer’s disease (AD), Parkinson’s disease (PD), frontotemporal lobar degeneration, and amyotrophic lateral sclerosis (ALS)^1^. To generate novel leads in tackling this growing problem, many genome-wide association studies (GWAS) have been performed in the various neurodegenerative diseases, involving >100,000 patients, and identifying >200 genetic risk loci^2^. While genetic risk loci have been utilized, singly or in aggregate, to refine predictions for risk of developing disease^3^,^4^, the greatest potential for these GWAS-identified loci may lie in the identification of novel disease mechanisms^5^.

However, the interpretation of disease-associated risk loci is complicated. The “sentinel” variant, usually a single nucleotide polymorphism (SNP) identified by GWAS, is rarely the specific change in DNA sequence – or “causal” variant – that results at the molecular level in a mechanistic change. Instead, in most cases, tens or hundreds of genetic variants at each locus are in strong linkage disequilibrium (LD) with the sentinel variant, constituting a set of co-inherited variants – or haplotype – any of which may be the underlying cause for increased disease risk^6^. Indeed, the risk-associated haplotype may span multiple genes, making even the gene to which a GWAS signal belongs unclear. Given these complexities, it is perhaps unsurprising that, with one exception pertaining to common variants near the *SNCA* gene, which was already implicated prior to the GWAS era in the development of PD^7^, none of the neurodegenerative disease risk loci identified by GWAS have been characterized in molecular detail. Yet such a molecularly precise understanding of a GWAS-identified genetic risk locus is a likely prerequisite for downstream therapeutic development.

Frontotemporal lobar degeneration (FTLD) is a neurodegenerative dementia affecting ~10-20 per 100,000 persons between the ages of 45 and 64, making FTLD the second most common early-onset dementia^8^,^9^. FTLD is a fatal, untreatable disease, with death typically occurring within ~8 years after diagnosis^8^. Noncoding single nucleotide polymorphisms (SNPs) on chromosome 7p21 have been associated with risk for the major neuropathological subtype of FTLD, characterized by pathological inclusions of the protein TDP-43 (FTLD-TDP)^10^. The association of this locus with FTLD-TDP has been replicated^11^^−^^13^, and the major T allele of the sentinel SNP, rs1990622, yielded an odds ratio of ~1.6 for disease development^10^. Genotype at rs1990622 also affects age at disease onset in Mendelian forms of FTLD-TDP^12^,^14^,^15^, as well as risk for development of cognitive impairment in the related disorder amyotrophic lateral sclerosis (ALS)^16^. We and others have implicated a gene in this region, *TMEM106B*, as causal^17^^−^^19^ However, studies to date have not explained how genetic variation at the 7p21 locus affects the function of *TMEM106B* or another gene, thereby contributing to the pathogenesis of FTLD-TDP.

In this study, we demonstrate that (1) common GWAS-implicated variants associated with FTLD-TDP are correlated with expression levels of *TMEM106B*, with increased expression correlating with the risk haplotype, (2) incremental increases in *TMEM106B* expression are associated with incremental increases in cell toxicity, (3) the risk allele of a candidate causal variant (rs1990620) in complete LD with rs1990622, the GWAS sentinel SNP, increases recruitment of the chromatin organizing protein CCCTC-binding factor (CTCF) downstream of *TMEM106B*, and (4) long-range chromatin looping interactions involving the CTCF site and other distal regulatory elements at the *TMEM106B* locus are stronger on the risk haplotype Together, these data provide a molecularly detailed mechanism for the effect of common genetic variation at this locus on risk for neurodegenerative disease.

## MATERIALS & METHODS

### eQTL analyses

The GWAS sentinel SNP, rs1990622^10^, was queried for association with all transcripts genome-wide using The Genotype-Tissue Expression (GTEx) eQTL database^20^, consisting of 7,051 samples and representing 44 different tissues from 449 healthy donors. GTEx eQTL plots were generated with *SNiPA*^21^. Conditional analyses and fine-mapping were performed using HapMap3-imputed genotypes from a published multi-ethnic LCL eQTL study^22^, as previously described^23^. In brief, gene expression data were normalized to the empirical average quantiles across all samples. Subsequently, the distribution of each gene expression trait was transformed to the quantiles of the standard normal distribution, separately within each population. The effects of known and unknown covariates were controlled for by principal component analysis. A *cis*-eQTL scan was performed by regressing the additive effect of each SNP within 1Mb of *TMEM106B* on gene expression by Bayesian regression, as implemented in SNPTEST^24^.

### Analysis of LD structure at the *TMEM106B* locus

HapMap Phase I, II and III combined CEU genotype data^25^ were visualized in HaploView to assign LD blocks^26^, filtering out SNPs with MAF<0.001 and requiring that 90% of informative pairwise LD values within a block must represent strong LD. Pairwise LD between variants at the *TMEM106B* locus was determined using both HapMap and 1000 Genomes data^27^, visualized either in HaploView or on the HaploReg v4.1 web resource^28^.

### Cell culture

HeLas were cultured in DMEM with 10% FBS, 1% L-glutamine, and 1% penicillin/streptomycin. LCLs were obtained from the Coriell Institute for Medical Research in Camden, NJ, and were cultured in RPMI with 15% FBS, 1% L-glutamine and 1% penicillin/streptomycin. Jurkats were cultured in RPMI with 10% FBS, 1% L-glutamine and 1% penicillin/streptomycin.

### *TMEM106B* overexpression experiments

*TMEM106B* expression constructs designed to over-express at 2-fold, 5-fold, and 20-fold expression, as previously described^19^, were transfected into HeLas with Lipofectamine 2000, as per manufacturer protocols. 48 hours post-transfection, ten brightfield images were taken across three biological replicates for each condition at 100X on a Life Technologies EVOS FL microscope. Image files were assigned random identifiers and a blinded individual counted the number of cells in each image that displayed the vacuolar phenotype, defined by having at least one clear punctate vacuolar structure. This experiment was repeated three times and the results were pooled. To assess cytotoxicity, the same transfection protocol was carried out, but at 48h post-transfection cells were spun down and resuspended in Trypan Blue-containing DMEM culture media, and the proportion of Trypan Blue-positive cells were determined with a hemocytometer. This experiment was also carried out three times. Western blots were performed for all six experiments as described previously^19^, and the data were pooled together for the quantification shown in **Figure 2B**.

### Cell line haplotype phasing

In order to confirm all cell lines used for mRNA stability experiments, ASE analysis, and Capture-C as *TMEM106B* haplotype heterozygotes, we first performed TaqMan SNP Genotyping assays, as previously described^15^, for rs1990622 and marker SNPs in strong or complete LD with rs1990622 (rs3807865, r^2^=0.9; rs6966915, r^2^=1; rs3173615, r^2^=1; rs1468803, r^2^=1), to confirm heterozygosity. For the cell lines used for the Capture-C experiments, the CTCF binding region was also analyzed by Sanger sequencing to confirm heterozygosity for the three completely linked potential causal variants: rs1990622, rs1990621, and rs1990620. Heterozygosity of the promoter SNP rs4721056 (r^2^=0.5 with rs1990622) was confirmed by genotyping as well, but because of the lower LD of this SNP with rs1990622, we PCR amplified the region containing rs4721056 and three SNPs in strong LD with rs1990622 (rs7781670, r^2^=0.9; rs1019309, r^2^=0.9; and rs1019307, r^2^=0.89) in the three Capture-C cell lines. This amplicon was cloned into the MCS of the pGL3 Promoter vector, and Sanger sequencing of individual clones confirmed that no cell lines had mixed haplotypes comprising these SNPs, thus linking the risk allele of rs4721056 to the risk haplotype in all cell lines.

### mRNA stability experiments

Three LCLs homozygous for the *TMEM106B* risk haplotype, and three LCLs homozygous for the protective haplotype, were treated with 1μg/μL Actinomycin D, and RNA was extracted at 0h, 1h, 2h, 4h, 8h and 24h post-treatment. RT-qPCR was performed as previously described^19^ to quantify *TMEM106B* levels at each time point, normalized to 18S RNA, which decayed by only ~14% in 24hr. Decay curves for mRNA stability were compared using two-way ANOVAs. This experiment was performed on each of the six cell lines twice, for a total of six biological replicates for each haplotype.

### Epigenomic prioritization of candidate *cis*-regulatory elements

We used the UCSC Genome Browser (GRCh37/hg19)^29^ to visualize ENCODE DNase hypersensitivity, transcription factor ChIP-seq, and the H3K4me1, H3K4me3, and H3K27ac histone mark tracks in the GM12878 LCL line^30^. We also visualized these histone marks in all LCLs included in the NIH Roadmap Epigenome Project ^31^ using the WashU Epigenome Browser^32^. First, we determined which of the top eQTL SNPs from the fine-mapping overlapped a *cis*-regulatory element (CRE) predicted to be active in LCLs, based on the presence of *at least one* of the features mentioned above. This permissive filtering process yielded only three candidate CREs with seven candidate causal variants. Two putative CREs displaying either DNase hypersensitivity, transcription factor binding, and/or H3K4me1 were tested for enhancer activity in LCL reporter assays. The third CRE contains three candidate causal variants and displays CTCF binding in LCLs, neuronal and glial cell lines, and was further investigated for allele-specific effects.

### Reporter assays

To test the two putative LCL CREs for enhancer activity, we cloned each region with either the risk or protective haplotype SNP alleles, based on 1000 Genomes Phase I^33^ haplotype information, into the upstream multiple cloning site (MCS) of the pGL3 Promoter luciferase reporter vector (Promega Cat. #E1761). We transfected 2.5x10^6^ LCLs using program Y-001 and nucleofection solution V on the Lonza Nucleofector 2b, using the pGL3 Basic vector (with no enhancer) (Promega Cat. #E1751) as a negative control, and the pGL3 Control vector (with an SV40 enhancer) (Promega Cat. #E1741) as a positive control (**Figure S3**). Three or four biological replicates were included for each construct in each experiment, and four independent experiments were performed for each candidate CRE. Cell lysates were isolated 24h post-transfection for luciferase readout using the Promega Dual-Luciferase Reporter Assay System (Promega Cat. #E1910), as described previously^19^. We used two-tailed t-tests to test for statistically significant differences in reporter activity.

### ENCODE data-mining for CTCF binding and DNase hypersensitivity

We downloaded the BAM read alignment files for all CTCF ChIP-seq experiments that showed a CTCF peak at the region containing rs1990620, as well as for the DNase digital genomic footprinting experiments performed in cell lines that show a DNase-seq peak at this region (https://www.encodeproject.org/, **Table S3)**. We analyzed raw sequencing reads containing rs1990620 to identify cell lines heterozygous at this locus. We summed risk allele reads across all cell types, as well as protective allele reads across all cell types, assessing for deviation from a 50:50 proportion using a two-tailed binomial sign test.

### Electrophoretic mobility shift assays

Nuclear extract was obtained from healthy human occipital cortex, LCLs, and HEK293s using the Thermo Fisher Scientific Nuclear and Cytoplasmic Extraction Reagents kit (Cat. #78833). A 61bp biotinylated DNA probe containing the risk or protective allele of rs1990620 at position 31 and 30bp of genomic sequence on either side was incubated with extract and competed with excess amounts of unlabeled oligo containing either the risk or protective allele of rs1990620. 2μL of anti-CTCF antibody (EMD Millipore Cat. #07-729) was added to the reaction to test for supershifts. EMSAs were performed in accordance with the Thermo Fisher Scientific LightShift Chemiluminescent EMSA kit (Cat. #20148).

### Capture-C

Capture-C was performed similarly to previous reports^34^. Briefly, 3C libraries were generated by fixing 10x10^6^ cells with formaldehyde, followed by *DpnII* digestion and ligation. Phenol/chloroform-extracted DNA was sonicated to produce 200-300bp fragments, and sequencing libraries were prepared with the NEBNext DNA Library Prep Master Mix Set (Illumina Cat. #E6040). 10μg of each capture library underwent multiplexed PCR with unique index oligos. Hybridization with 60bp biotinylated capture probes (**Table S4**) was performed with the SeqCap EZ Hybridization and Wash Kit (Roche Cat. #05634261001). Briefly, 3C libraries were air-dried with heat in a thermocycler, and resuspended with hybridization reagents. 2μL (3pmoles) of pooled capture probes for each bait region were added to the resuspended libraries, and incubated for 72h in a thermocycler at 47 degrees C. After isolation of captured material using streptavidin beads and PCR, an additional 24h capture was performed. The capture probes, ordered from Integrated DNA Technologies, flank *DpnII* cut sites that are proximal to marker SNPs in the *TMEM106B* promoter region and CTCF site, designed for captured ligation products to contain marker SNPs distinguishing between haplotypes. Promoter capture experiments used rs4721056 to distinguish haplotypes, and CTCF site capture experiments used rs1990620 and rs1990621 to distinguish haplotypes (see “Cell line haplotype phasing” section). Samples were pooled and sequenced on one lane of an Illumina HiSeq 2500 with 125bp paired end reads, yielding ~230 million read pairs. Two LCLs and Jurkat cells were captured at both regions, with two technical replicates for each capture, performed by different individuals in tandem.

### Capture-C data analyses

Quality control was performed with FastQC (http://www.bioinformatics.babraham.ac.uk/projects/fastqc/) and pre-processing and read alignment were performed as previously described^35^. Briefly, the paired-end reads were reconstructed into single reads using FLASH, *in silico*-digested using the DpnII2E.pl script (https://github.com/telenius/captureC/releases), and mapped with Bowtie1^36^. The aligned reads were then analyzed with the CCanalyser3.pl script (https://github.com/telenius/captureC/releases). Interactions were tested for significance with fourSig^37^, using default parameters and a window size of five. First, we tested interactions for significance using all reads mapping to chromosome 7. Then we restricted our analyses to interactions within the TAD and sub-TAD to test for significant interactions *within* these regions, respectively.

To determine if the long-range interactions captured by the hybridization probes show allelic bias, we first estimated the technical (non-biological) bias for each SNP, which may reflect capture bias, mapping bias, or other sources of technical bias. We estimated technical bias using two independent approaches. For the first approach, we enumerated the risk vs. protective allele reads containing each marker SNP directly *adjacent* to the probe sequences (*i.e.* mapping to the 5kb window containing the probe sequences or the immediately adjacent 5kb windows at each capture site) for each experiment. In the second approach, we enumerated the risk vs. protective allele reads containing each marker SNP mapping *at a great distance* from *TMEM106B* (*i.e.* outside of the *TMEM106B* TAD, as defined by the LCL Hi-C heatmap^38^). In each case, we assumed that the interactions too close or too far from *TMEM106B* do not represent functional interactions. For the adjacent interactions, ligations may occur artifactually due to chromosomal proximity^39^. For the interactions outside the *TMEM106B* TAD, low read count/interaction suggests that these are not true interactions, consistent with the fourSig results. Technical bias estimated by these two approaches is likely conservative – some true biological interactions may be “discounted” in this way, creating false negatives in order to most reliably account for false positive interactions. Despite different underlying assumptions, technical bias estimates derived from the two methods agree with each other to within <1% for both promoter and CTCF site marker SNPs.

In each case, read counts for interactions originating from each haplotype were compared (1) with an expected proportion of 0.5 (in analyses without normalization for technical bias) or (2) with the proportions observed using the estimates of technical bias described above (in analyses normalized for technical bias), using a binomial test.

### CTCF site SNP enrichment analysis

We identified all SNPs that have been associated with risk for FTLD, Alzheimer’s disease, Parkinson’s disease, amyotrophic lateral sclerosis, and related conditions, at a genomewide significant level (p≤5x10^−8^), using the EMBL-EBI/NHGRI GWAS catalog^2^. This list contains 200 SNPs associated with 29 traits. We downloaded all available optimal irreproducible discovery rate (IDR) thresholded peak^40^ BED files from ENCODE CTCF ChIP-seq experiments performed in brain-relevant cell/tissue types (https://www.encodeproject.org/; hippocampal astrocytes, cerebellar astrocytes, BE2_C neuroblastoma, retinoic acid-treated SK-N-SH neuroblastoma, choroid plexus epithelial cells, H54 glioblastoma, and brain microvascular endothelial cells). We used the GREGOR pipeline^41^ to determine if the disease-associated SNPs or their LD proxies (at r^2^ values of >0.7, >0.8 and >0.9) were enriched in brain CTCF binding sites compared to control SNPs that were matched for minor allele frequency, number of LD proxies, and distance to the nearest gene^41^. 14 of the neurodegenerative risk SNPs were not identifiable by GREGOR, resulting in a final list of 186 disease-associated SNPs and proxies.

## RESULTS

### Genetic variation at the 7p21 locus associates with TMEM106B expression

It is increasingly recognized that many GWAS-implicated variants associated with disease risk may confer their effects by altering the expression levels of nearby genes^7,42-49^. In the case of 7p21, several studies have demonstrated such an expression quantitative trait locus (eQTL) effect for *TMEM106B* in multiple human tissue types, including brain and Epstein-Barr virus (EBV)-immortalized B lymphoblastoid cell lines (LCLs)^22^,^50^^−^^52^. We therefore systematically investigated the 7p21 locus for all eQTL effects, in order to confirm the *TMEM106B* eQTL effect and to exclude other potential causal genes at this locus.

Analysis of data from The Genotype-Tissue Expression (GTEx) Project^20^ demonstrated a robust association between genotype at rs1990622, the GWAS sentinel SNP, and *TMEM106B* expression in several cell types from normal individuals (**Figure 1A**), including LCLs (n=114, *P*=1.9x10^−6^) and transformed fibroblasts (n=272, *P*=3.0x10^−7^). No other transcript genome-wide was significantly associated with rs1990622 genotype, consistent with other published large-scale eQTL studies^50^,^51^. The association between rs1990622 and *TMEM106B* levels has also been previously reported in human brain^52^; GTEx data confirms this relationship in the hippocampal and nucleus accumbens brain regions (**Figure 1B**). In all cell types, the risk allele was consistently associated with increased *TMEM106B* expression.

**Figure 1.**
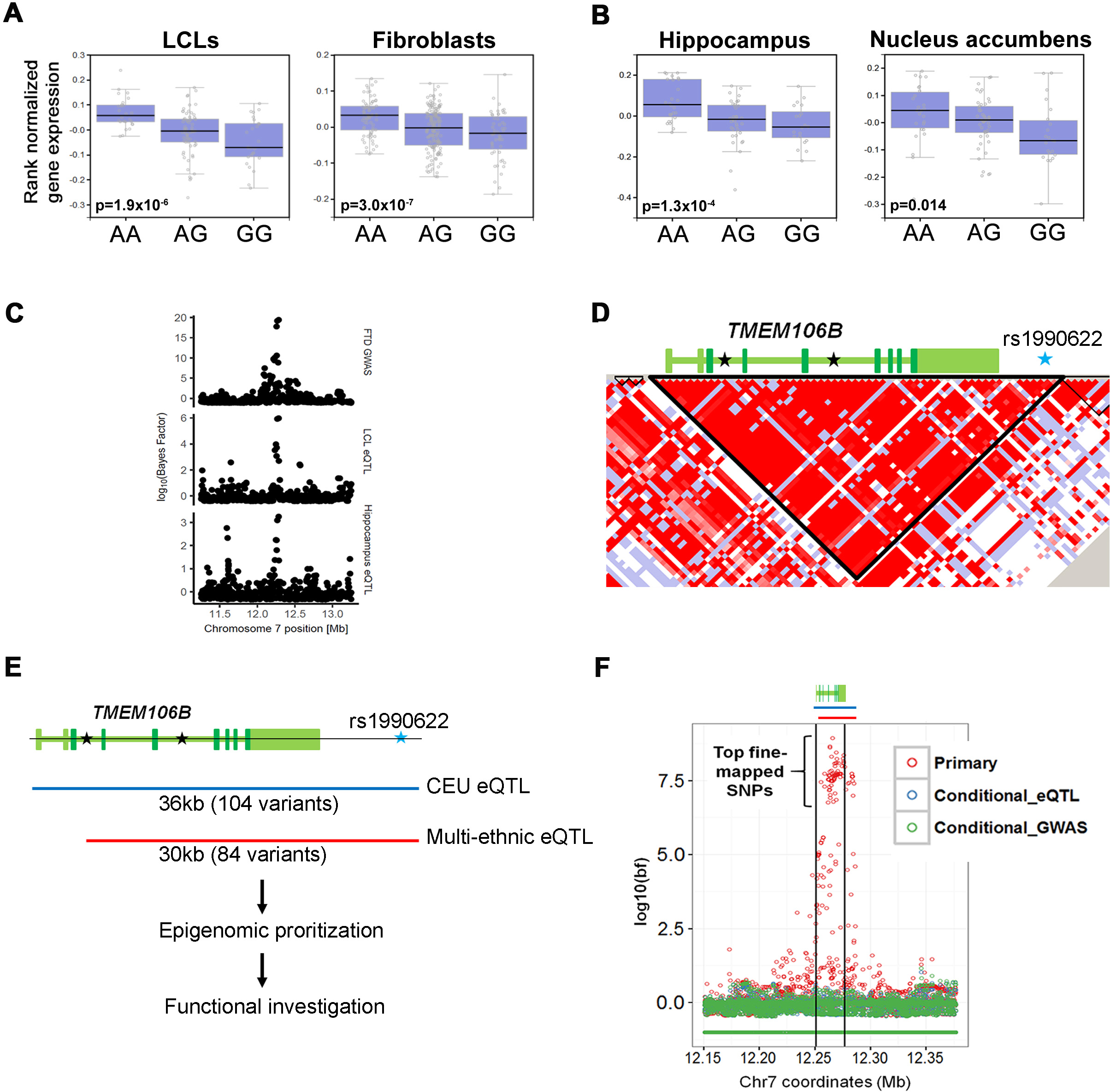
Analysis of expression quantitative trait locus (eQTL) effect at *TMEM106B*. Boxplots from the Genotype-Tissue Expression (GTEx) Project data demonstrate association of *TMEM106B* levels with rs1990622 genotype (A=risk allele) in peripheral cell types **(A)** as well as human brain regions **(B)**. Data from LCLs (n=114), fibroblasts (n=272), hippocampus (n=81) and nucleus accumbens (n=93) are shown. Black lines indicate median expression, lower and upper bounds of boxes indicate 25^th^ and 75^th^ percentile expression levels, respectively, and circles outside whiskers denote outliers. **(C)** Colocalization analyses over a 2Mb region centered on *TMEM106B*. Both the LCL (middle row) and hippocampal (bottom row) eQTL association signals overlap the FTLD-TDP risk association signal (top row). Each circle represents a SNP, with genomic position on the x-axis and significance of association on the y-axis (log10- transformed Bayes factor). (**D, E**) LD structure at *TMEM106B* in CEU populations, with gene structure indicated above LD plot. Coding exons are in dark green, UTRs are in light green, and SNPs associated by GWAS with frontotemporal lobar degeneration (FTLD) are indicated with stars, including the sentinel SNP, rs1990622. The *TMEM106B* eQTL effect extends across the 36kb LD block, but can be truncated on the 5’ end with the addition of ethnicities with different haplotype structures **(E)**. **(F)** Multi-ethnic conditional eQTL analysis of *TMEM106B* locus. Each circle represents a SNP, with genomic position on the x-axis and association with *TMEM106B* levels on the y-axis (log_10_-transformed Bayes factor). *TMEM106B* gene and regions of eQTL association are indicated above the plot. The primary multi-ethnic analysis (red) demonstrates a strong association between a SNP cluster and *TMEM106B* expression. Conditioning this analysis on either the top eQTL SNP (blue) or the sentinel GWAS SNP (green) results in loss of an association signal at this locus (*i.e.* there are no highly-associated SNPs shown in blue or green), suggesting that there is only one eQTL signal and that the causal variant underlying association with *TMEM106B* expression may be the same as the causal variant underlying association with FTLD.

In order to determine whether these eQTL signals are likely to result from the same functional genetic variant(s) underlying risk for FTLD-TDP, we performed colocalization analyses over a 2Mb region centered on *TMEM106B*. Both the LCL and hippocampal eQTL association signals overlap the FTLD-TDP risk association signal (**Figure 1C**). Specifically, the LCL eQTL signal has a 97% posterior probability of representing the same signal as the association with risk for FTLD-TDP, thus making LCLs an attractive cellular model to investigate the molecular underpinnings of disease association at this locus.

The 7p21/*TMEM106B* locus is harbored within a 36kb LD block in samples from individuals of European ancestry, the population in which the original FTLD-TDP GWAS was performed (**Figure 1D**). This LD block encompasses the *TMEM106B* promoter, the entirety of the *TMEM106B* gene, and extends ~10kb downstream of the gene. According to 1000 Genomes data^53^, the block contains 104 genetic variants that are in strong, but not perfect, LD with rs1990622 (r^2^>0.8, **Figure 1D**). Indeed, in-depth examination of the eQTL effect in human hippocampal or LCL (**Figure S1**) samples from GTEx reveals dozens of variants in strong LD with rs1990622 that are associated with *TMEM106B* expression to a similar degree. We thus asked whether more than one eQTL signal occurs in this region, and what the candidate causal variant(s) underlying association with disease or expression might be.

We began by honing the region of eQTL association. To do this, we performed a second eQTL analysis of LCLs from eight ethnic populations^22^, reasoning that the different haplotype structures seen in disparate populations might refine the 36kb LD block of association seen in individuals of European ancestry. We found that with the addition of these populations, the region of association with *TMEM106B* expression could be truncated on the 5’ end, effectively removing the promoter and first two exons of the gene, and reducing the number of potential causal variants to 84 (75 SNPs and 9 indels, **Figure 1E**).

We then performed conditional analyses using the refined region of association from the multi-ethnic analysis^22^. Conditioning on either the GWAS sentinel SNP, rs1990622, or the most significant eQTL SNP, rs6948844 (r^2^=1 with rs1990622), yielded no variants within a 2Mb region that demonstrated any residual association with *TMEM106B* expression (**Figure 1F**) These results suggest that there is only one eQTL signal at this locus, and that the causal variant underlying the association with *TMEM106B* expression may be the same as the causal variant underlying association with disease.

### Increased levels of TMEM106B expression correlate with increased cellular toxicity

If the causal variant responsible for association with *TMEM106B* expression levels confers risk for neurodegeneration, one would expect incremental changes in *TMEM106B* expression to lead to incremental effects on cellular health. We and others have previously shown that over-expression of *TMEM106B* results in the development of enlarged LAMP1+ late endosomes/lysosomes appearing as vacuolar structures in multiple cell types, including neurons, with associated impairment in lysosomal degradative function^18^,^19^,^54^,^55^. However, the magnitudes of reported eQTL effects in human tissues are often modest ^20^,^56^, and so we sought to understand the effects of incremental increases in *TMEM106B* expression on disease-relevant measures such as (1) development of the previously-reported vacuolar phenotype and (2) cell toxicity.

To do this, we employed three different *TMEM106B* constructs that reliably produced a spectrum of over-expression ranging from ~2X to ~20X (**Figures 2A and 2B**). In HeLa cells, we found that with each incremental increase in *TMEM106B* expression over baseline, the percentage of cells exhibiting the vacuolar phenotype of enlarged lysosomes increased (**Figures 2C and 2D**), along with increased cell death (**Figure 2E**). Together, these findings suggest that genetic variation at the 7p21 locus may influence risk for neurodegeneration by altering *TMEM106B* expression-dependent effects on lysosomal function and cell health.

**Figure 2.**
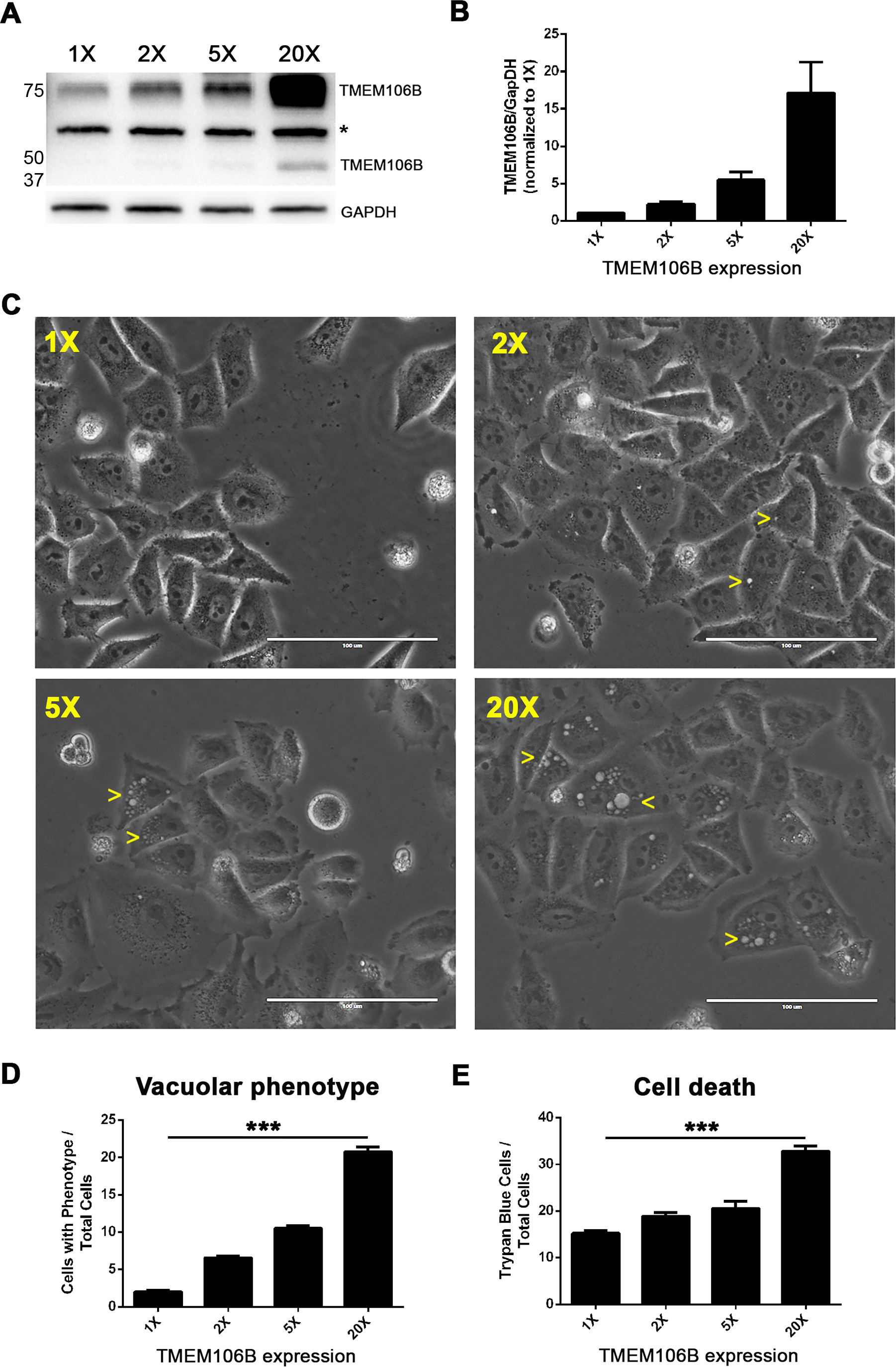
TMEM106B expression levels show dose-dependent effects on cell toxicity. **(A)** Representative Western blot of TMEM106B levels in the absence (1X) and presence (2X, 5X, 20X) of various *TMEM106B* over-expression constructs transfected into HeLas. The bands at ~75kD and ~40kD represent dimeric and monomeric forms of TMEM106B. A non-specific band is indicated by the asterisk. **(B)** Quantification of blots from six independent experiments (+/- SEM) demonstrate reliable expression levels of each construct. **(C)** Representative bright-field images demonstrate a dose-dependent vacuolar phenotype in cells. Yellow arrowheads indicate cells exhibiting the phenotype. Quantification of the number of cells exhibiting **(D)** the vacuolar phenotype and **(E)** cell death is shown for each expression paradigm, across three independent experiments. Asterisks denote statistical significance (p<0.001 by ANOVA).

### A candidate causal regulatory region

We next sought to identify the causal variant or variants responsible for allele-specific regulation of *TMEM106B* expression and, by extension, risk for FTLD-TDP. *TMEM106B* steady-state transcript levels depend on both the production of mRNA and its stability. We first considered the possibility of differential mRNA stability. In multiple LCLs homozygous at the *TMEM106B* locus, we found that mRNA stability did not differ between risk haplotype homozygotes and protective haplotype homozygotes (**Figure S2**), suggesting that differences in the production of mRNA account for the observed eQTL effect.

To identify variants that may have transcriptional regulatory effects, we examined the 84 candidate variants (75 SNPs and 9 indels) located within the refined region of eQTL association (**Figures 1E and 1F**). We used ENCODE^30^ and NIH Roadmap Epigenome^31^ data to determine whether each variant is located in a predicted LCL *cis-*regulatory element (CRE), as determined by DNase I hypersensitivity (DHS), transcription factor (TF) binding, or the active histone marks H3K27ac, H3K4me1 or H3K4me3.

Surprisingly, only seven SNPs spanning three candidate regulatory regions met these permissive conditions (**Figure 3 and Tables S1 and S2)**. Three SNPs located in intron 4 of *TMEM106B* overlap a region of DHS, TF binding, and the enhancer-associated histone mark H3K4me1 in LCLs, while a fourth SNP downstream of *TMEM106B* overlaps a binding site for the TF PU.1 in LCLs. Upon empirical testing in luciferase reporter assays, however, these regions displayed little or no enhancer activity in LCLs, consistent with the lack of the H3K27ac histone mark, which has been reported to distinguish active from inactive or poised enhancers ^57^,^58^. Furthermore, the risk and protective haplotype versions of these regions did not differ in activity, suggesting that none of the overlapping SNPs affect regulatory activity (**Figure S3**). The remaining candidate regulatory region contains the remaining three completely linked SNPs, one of which is rs1990622, the GWAS sentinel SNP. Moreover, in virtually all ENCODE-tested cell types, including neuronal and glial lines, this putative CRE displays binding for the mammalian chromatin organizing protein CTCF. Therefore, we investigated this region for potential allele-specific effects (**Figure 3**).

**Figure 3.**
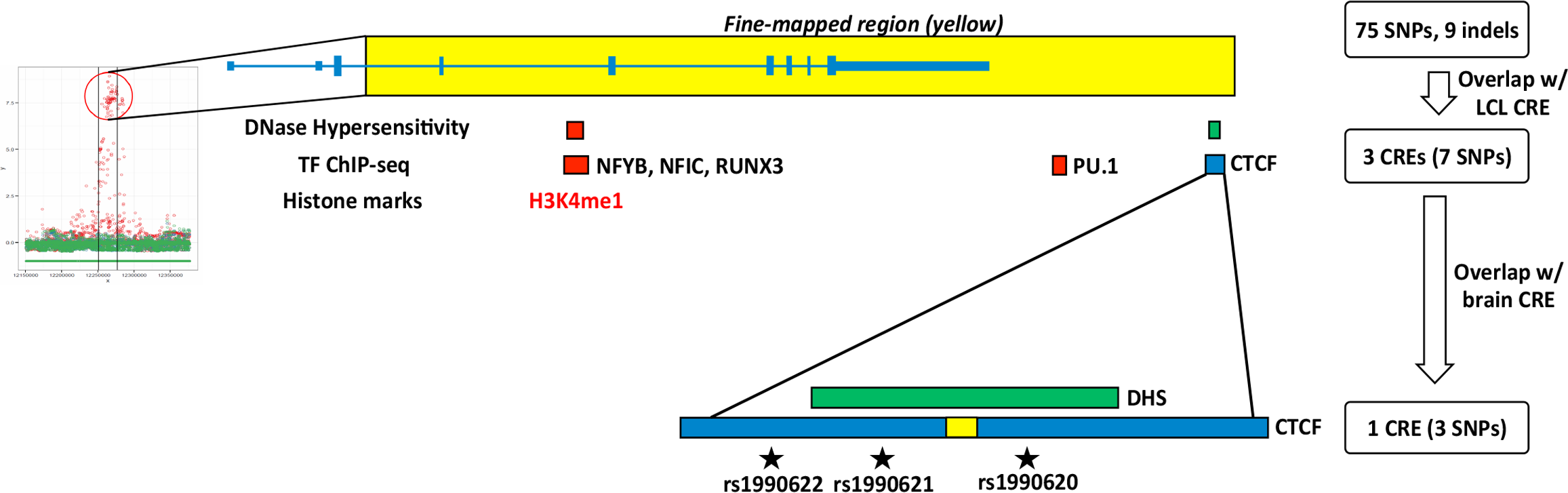
Prioritization of *cis*-regulatory elements (CREs) harboring candidate functional variants. The 84 variants from the eQTL fine-mapping (left) were prioritized based on overlap with predicted CREs in LCLs (red boxes and text), neuronal and glial cell lines (green), or all three (blue), based on ENCODE and NIH Roadmap Epigenome project data (see flow chart at right of image). This analysis yields 7 SNPs in 3 potential CREs as candidate causal variants. Only one CRE – an intergenic CTCF binding site (CTCF motif represented as yellow rectangle) – is predicted to be active in brain tissue; this CTCF-binding CRE contains three SNPs in complete LD, including the GWAS sentinel SNP, rs1990622.

### Common variation at rs1990620 affects binding of CTCF at the TMEM106B locus

Given the observed eQTL effect at *TMEM106B* in multiple tissues, and the CTCF binding site within this potential CRE, we next sought evidence for allele-specific binding of CTCF to our candidate regulatory region. To do this, we analyzed ENCODE CTCF ChIP-seq and DNase digital genomic footprinting (DGF) experiments. CTCF ChIP-seq was performed on 99 ENCODE cell lines, 95 of which show a CTCF peak at the putative CRE. We were able to confirm 20 of these cell lines as *TMEM106B* haplotype heterozygotes by examining reads containing rs1990620, the SNP closest to the CTCF core motif (48bp from core motif) and covered by the most reads (**Figure 4A and Table S3)**. By analyzing the reads covering rs1990620, we found significant enrichment of CTCF binding to the risk-associated A allele (*P*=0.043, **Figure 4B**). In addition, we identified 6 cell lines heterozygous at rs1990620 that were interrogated by DNase DGF. In these lines, we found that the chromosome bearing the risk A allele was significantly more sensitive to DNase cleavage (*P*<0.001, **Figure 4B**), consistent with an open chromatin state and potential regulatory activity^59^.

**Figure 4.**
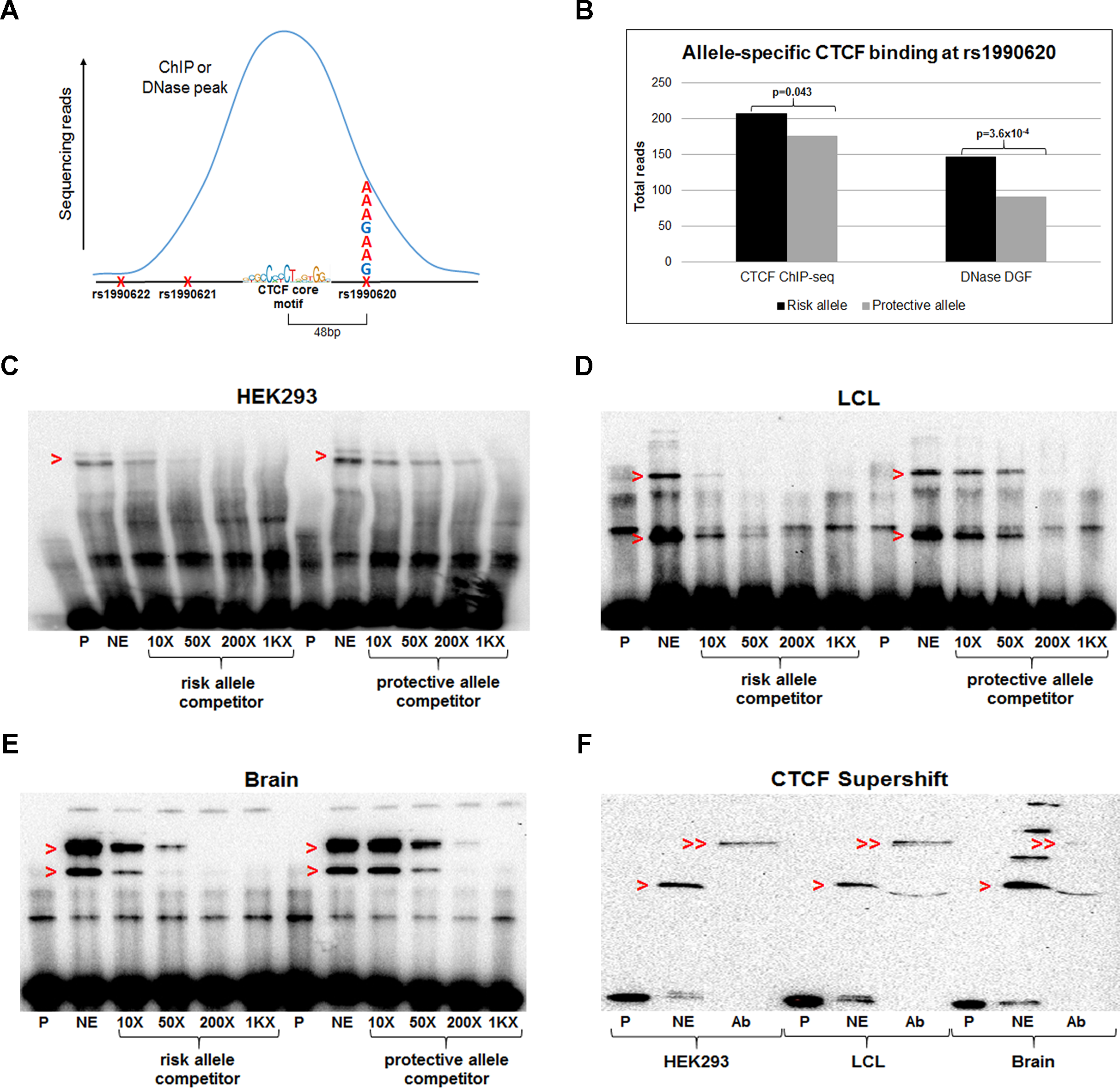
The risk allele of rs1990620 preferentially recruits CTCF in human tissues. **(A)** Schematic of approach for determining allelic bias in CTCF ChIP-seq and DNase digital genomic footprinting (DGF) experiments. The rs1990620 SNP (48bp from the CTCF core motif) was analyzed for number of reads containing the risk vs. protective allele in samples showing a CTCF ChIP-seq or DNase DGF peak at this region. **(B)** The risk allele of rs1990620 increases CTCF binding and DNase hypersensitivity at this region, based on data from 20 and 6 cell types heterozygous at this locus, respectively (**Table S3**). **(C-E)** A 5’ biotinylated probe (P) containing the rs1990620 risk allele was incubated with nuclear extract (NE) from HEK293s **(C)**, LCLs **(D)**, and human brain **(E)**. In each extract, the shifted probe/protein complex (red arrowheads) was more efficiently competed with an unlabeled competitor oligonucleotide (at 10X, 50X, 200X, or 1000X (1KX) the concentration of the labeled probe) containing the risk allele vs. the protective allele, indicating preferential binding of a nuclear factor to the risk allele. **(F)** A 3’ biotinylated probe (P) containing the rs1990620 risk allele is shifted by addition of nuclear extract (NE) from HEK293s, LCLs and brain (red arrowheads). Addition of an anti-CTCF antibody (Ab) results in disappearance of the shift and a supershift (double arrowheads), indicating that the complex binding to the probe contains CTCF.

We corroborated these data with *in vitro* investigations of CTCF binding by electromobility shift assays (EMSA). Utilizing a competition EMSA approach, we found that the risk allele of rs1990620 was indeed more effective at shifting a protein complex in extracts from HEK293 cells (**Figure 4C**), LCLs, (**Figure 4D**), and human brain (**Figures 4E and S4**). Addition of CTCF antibody resulted in both a supershift and disappearance of the original EMSA band, demonstrating that the complex preferentially binding to the A allele of rs1990620 contains CTCF (**Figure 4F**). Thus, ChIP-seq-based investigation of differential CTCF binding by allele, coupled with gel shift assays, together indicate that common variation at a single SNP, rs1990620, may underlie haplotype-specific effects on CTCF recruitment.

### Long-range interactions involving TMEM106B demonstrate haplotype-specific effects

Rapidly emerging evidence suggests that CTCF plays a major role in the shaping of the three-dimensional architecture of the mammalian genome^60^,^61^. In particular, CTCF has been reported to contribute to the formation of topologically associated domains (TADs), which may be central to enhancer-promoter interactions and insulator function ^62^^−^^66^. Indeed, interrogation of high resolution *in situ* Hi-C data from LCLs^38^ suggests that the CTCF binding region containing rs1990620 is located within a small ~250kb TAD (sub-TAD) that is part of a larger ~1Mb TAD, and is involved in multiple long-range looping interactions (**Figure S5)**. Moreover, one of the major interactions occurs between this region and the *TMEM106B* promoter, implicating a role for this CTCF site in *TMEM106B* regulation (**Figure 5A).** Interestingly, there are also strong Hi-C interactions between the promoter and a predicted enhancer of *TMEM106B*^67^ located ~13kb downstream of the CTCF site. This enhancer also lies within the sub-TAD and harbors no disease-associated genetic variants. Given the observed allele-specific effects on CTCF recruitment, we hypothesized that rs1990620 may influence *TMEM106B* expression and, by extension, neurodegenerative disease risk, through differential effects on CTCF-mediated interactions between distal regulatory elements ^68^ (**Figure 5B**).

**Figure 5:**
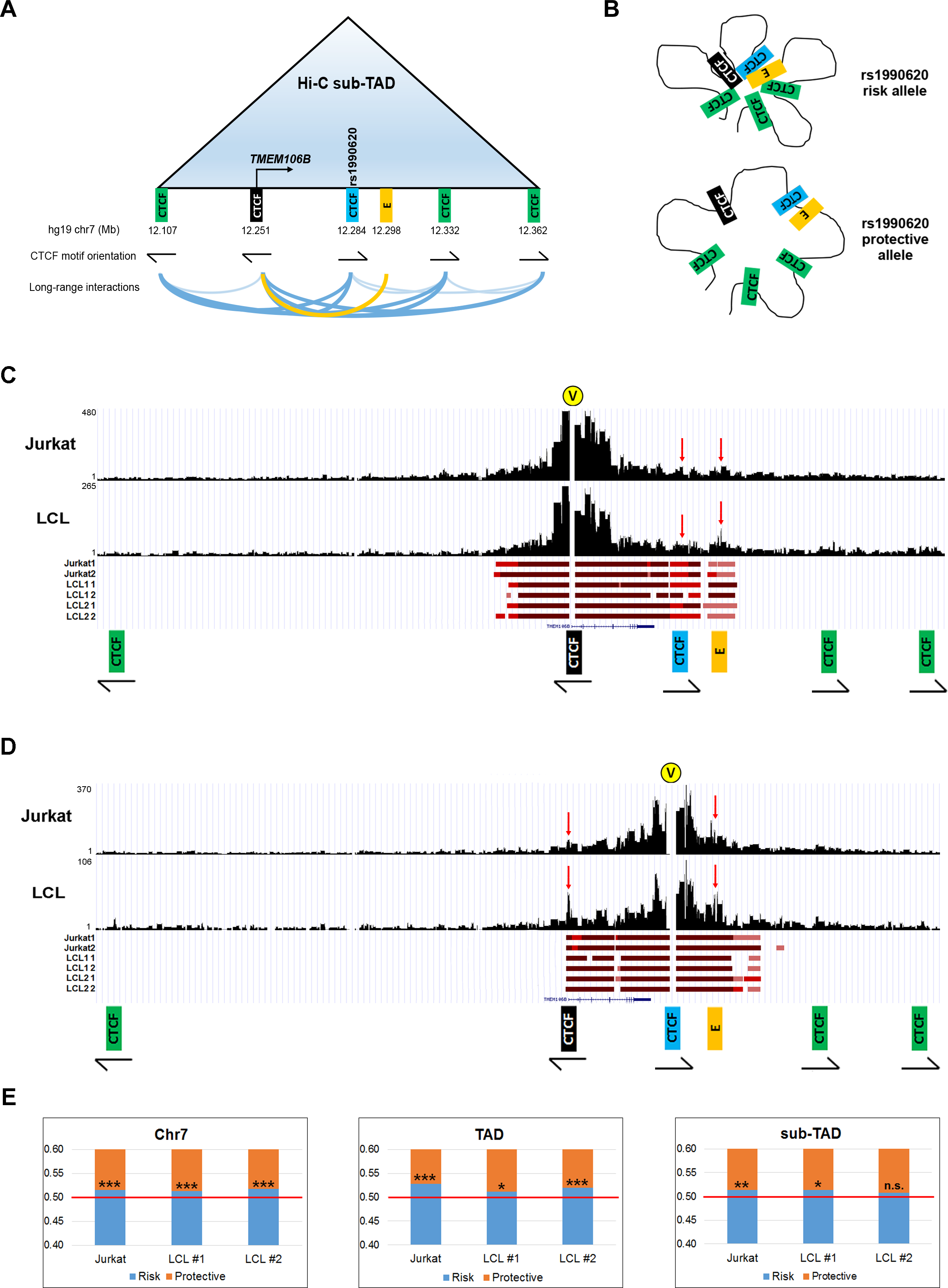
Haplotype-specific long-range chromatin interactions at the *TMEM106B* locus. **(A)** Schematic representation of the *TMEM106B* sub-TAD and interactions among distal regulatory elements, based on published LCL Hi-C data. The black CTCF site is located at the *TMEM106B* promoter; the blue CTCF site contains rs1990620; the gold rectangle labeled “E” represents a transcriptionally active enhancer. Note that the CTCF motifs present at the sub-TAD boundaries (12.107 and 12.362) follow the convergent orientation (arrows indicate direction and strand) most commonly reported for interacting CTCF sites. Darker lines in bottom of panel indicate interactions between CTCF sites in convergent orientation. **(B)** Model illustrating how allele-specific CTCF binding at rs1990620 might affect sub-TAD structure and long-range interactions at this locus, with more contact among distal regulatory elements bearing the risk-associated haplotype. **(C, D)** Capture-C experimental data for representative Jurkat and LCL samples, with raw read coverage shown on the y-axis for interactions captured by probes for (**C**) the *TMEM106B* promoter and **(D)** the rs1990620-containing CTCF site. Significant interactions within the sub-TAD for each cell line and replicate (3 cell lines with 2 replicates each) are indicated with red bars, below the coverage plots, with darker shades of red indicating higher confidence interactions. Yellow circles marked “V” indicate viewpoints (capture sites). Red arrows indicate interactions between the promoter, the rs1990620 CTCF site, and enhancer. **(E)** Allelic bias in long-range interactions involving the *TMEM106B* promoter across all of chromosome 7 (left), the 1Mb TAD (middle), and the 250kb sub-TAD (right) containing *TMEM106B*. Read count proportions from capture experiments containing either the risk (blue) or protective (orange) allele of a marker SNP are shown; in each case, more interactions with the *TMEM106B* promoter involve the risk haplotype. *p<0.01; **p<0.001; ***p<0.0001; n. s.=non-significant.

To test this, we adapted a recently-developed variation of chromosome conformation capture (Capture-C)^34^ to agnostically capture all interactions involving the CTCF-binding region, as well as the *TMEM106B* promoter. Specifically, we coupled 3C library preparation with a probe capture step to enrich for interactions involving our two regions of interest (**Table S4**). Importantly, we designed our capture probes not to overlap SNPs (thus, giving probes equal opportunity to bind to either allele), while also localizing to regions within 60bp of one or more marker SNPs (thus allowing for analysis of captured interactions in a haplotype-specific manner). We performed our Capture-C experiments in three different cell lines – two different LCLs and the T cell leukemia-derived Jurkat cell line (the *TMEM106B* eQTL effect has also been reported in T cells^69^,^70^) – with all three lines heterozygous for the *TMEM106B* haplotype.

When analyzing all long-range (≥2kb) interactions mapping to chromosome 7, we found that statistically significant interactions (based on an FDR threshold of 1% using fourSig^37^) were largely centered around the ~1Mb TAD containing *TMEM106B* (**Figure S6**). When restricting the application of fourSig to regions *within* the TAD (thus increasing the background model), all statistically significant interactions mapped to the sub-TAD (**Figure S7**). Thus, our data confirm the hierarchical nature of the chromatin architecture previously reported at this locus by *in situ* Hi-C. Importantly, all pair-wise interactions between the five sub-TAD CTCF sites, including the *TMEM106B* promoter, and the enhancer, were statistically significant in every sample, under both analyses (**Figure S7**), with the outermost CTCF sites in convergent orientation and delineating the boundaries of a topological domain.

To obtain a finer-grained understanding of the most meaningful interactions at our locus, we further restricted the fourSig analysis to the ~250kb sub-TAD, which further increased the background threshold for significance. Under these conditions, significant interactions between the *TMEM106B* promoter, the rs1990620-containing CTCF site, and the enhancer emerged (**Figures 5C and 5D**). These results implicate the CTCF site and the enhancer as potential key regulators of *TMEM106B*.

Recent studies suggest that genes involved in CTCF-associated long-range interactions tend to be more transcriptionally active than genes not involved in such interactions^38^,^68^. Therefore, we asked whether the number of long-range chromatin looping interactions involving the risk haplotype, which preferentially binds CTCF and expresses *TMEM106B* at higher levels, is significantly higher than the interactions involving the protective haplotype, in these heterozygous cell lines (**Figure 5A**). In all three cell lines, we observed significantly more interactions captured with the promoter probes occurring on the risk haplotype, with consistent effects whether we analyzed all interactions mapping to chromosome 7, or restricted the analysis to interactions within the TAD or sub-TAD (**Figure 5E**). When interactions were captured with probes targeting the rs1990620-containing CTCF binding site, fewer reads were obtained, and no apparent difference in raw reads involving the risk vs. protective haplotype were detected.

Given that various sources of technical bias (*e.g.* capture bias, alignment bias, bias in duplicate removal) can influence allele-specific high-throughput sequencing analyses, we next compared the number of long-range interactions involving each haplotype (“true” interactions) against the number of reads from each haplotype aligning to regions directly adjacent to the bait regions (“false” interactions,). We assumed that regions directly adjacent to the bait regions are subject to artefactual ligations due simply to chromosomal proximity, as has been previously suggested^39^. While some true functional interactions may be lost in this way (creating false negatives), this approach let us test for false-positive differences in interactions between the two haplotypes by capturing biases in read alignment or other technical steps of the Capture-C protocol.

After adjustment for technical bias, we still observed significant enrichment of promoter-captured interactions on the risk haplotype in two out of three cell lines (*P*=1.98x10^−2^ for Jurkats and *P*=5.78×10^−3^ for LCL Line 2 for significant deviation from expected proportions). Moreover, the same two cell lines demonstrated a significant enrichment of CTCF-associated interactions on the risk haplotype as well, whether probes captured interactions adjacent to the rs1990620 SNP (*P*=3.28x10^−3^ for Jurkats and *P*=4.37x10^−3^ for LCL Line 2) or the rs1990621 SNP (*P*<1.00x10^−6^ for Jurkats and *P*=2.38x10^−2^ for LCL Line 2, see **Figures 3** and **4A** for SNP locations).

Taken together, these data suggest that SNP-specific effects on CTCF recruitment may alter the genomic architecture at the *TMEM106B* locus, with ensuing alterations in gene expression.

### Common genetic variants associated with neurodegenerative diseases are enriched in brain CTCF binding sites

CTCF is emerging as a master regulator of mammalian gene expression through its widespread influences on genomic architecture^60^,^61^,^71^. Having uncovered an allele-specific effect on the expression of *TMEM106B,* a frontotemporal dementia-associated genetic risk factor, likely mediated by CTCF, we hypothesized that CTCF-mediated effects may play a more general role in conferring risk for neurodegeneration.

To test this, we identified all published SNPs associated at a genome-wide statistical significance level with risk for four major neurodegenerative diseases (FTLD, AD, PD, and ALS). 200 neurodegenerative disease risk SNPs were identified from the GWAS Catalog^2^, and this set of SNPs, as well as their LD proxies, were investigated for the extent of overlap with brain CTCF binding sites identified by ChIP-seq^30^.

Seven human brain-relevant CTCF ChIP-seq datasets were identified from ENCODE project data^30^. Across the combined set of all seven datasets, using the Genomic Regulatory Elements and GWAS Overlap algorithm (GREGOR^41^), we found a highly-significant ~1.5-fold enrichment of neurodegenerative disease SNPs and their LD proxies overlapping CTCF binding sites (*P*<0.0001 at an LD threshold of r^2^>0.7, **Figure 6A**). Evaluating each ChIP-seq dataset individually, CTCF binding sites found in brain-derived microvascular endothelial cells (1.6-fold enrichment, *P*<0.001), cerebellar astrocytes (1.4-fold enrichment, *P*=0.006), and neuroblastoma cells (1.5-fold enrichment, *P*=0.008) were most highly enriched for the presence of neurodegenerative risk SNPs, whereas CTCF binding sites found in other brain-derived cell types, including choroid plexus epithelial cells, showed no significant enrichment (**Figure 6B**). Our results are thus compatible with a wider role for CTCF-mediated effects in modulating risk for neurodegeneration.

**Figure 6:**
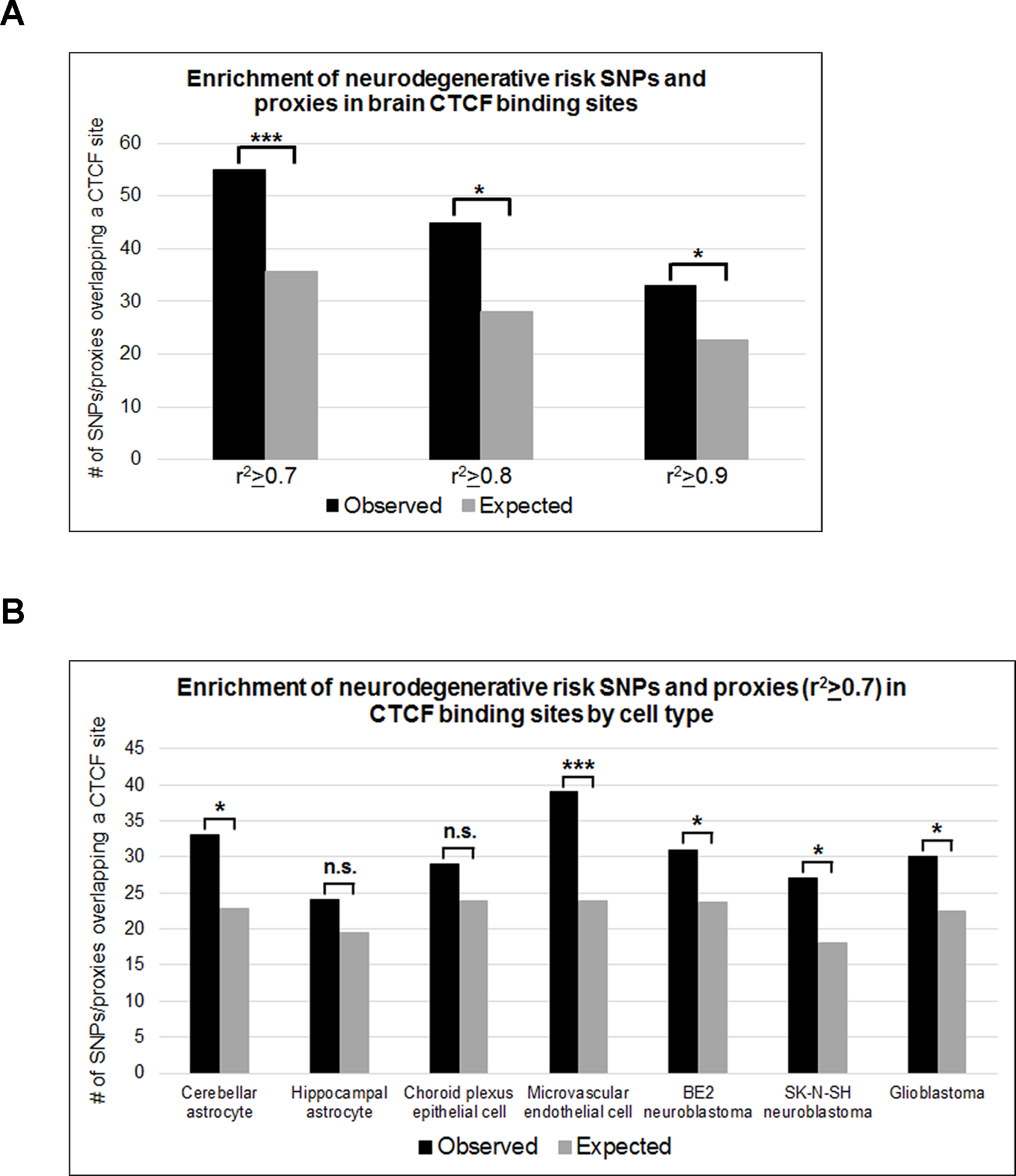
Neurodegenerative disease risk SNPs are enriched in brain CTCF binding sites. We determined the overlap between all SNPs identified by GWAS to confer risk for four different neurodegenerative diseases, as well as their proxies (at LD thresholds of r^2^≥0.7, ≥0.8 or ≥0.9), with brain CTCF binding sites. We then compared this “observed” overlap with an “expected” overlap with brain CTCF binding sites, based on 500 control SNPs matched to each GWAS SNP for minor allele frequency, number of LD proxies at each threshold, and distance to nearest gene, using the GREGOR algorithm. Statistically significant enrichments of neurodegenerative disease risk SNPs at CTCF binding sites were seen in the combined data sets at each LD threshold for proxy determination **(A)**, and in 5 out of 7 brain-relevant cell types, shown at an LD threshold of r^2^≥0.7 **(B)**. *p<0.05; **p<0.005; ***p<0.0005; n.s.=non-significant.

## DISCUSSION

Here, we functionally dissect a locus first linked to the human neurodegenerative disease FTLD by GWAS. Through a combination of data-mining and bench-based experimental studies of human-derived tissues, we demonstrate that common variants linked to FTLD by GWAS associate with haplotype-specific expression of *TMEM106B* in multiple tissues including human brain; that this effect may depend on haplotype-specific effects on recruitment of CTCF, with corresponding haplotype-specific effects on long-range chromatin interactions; and that incremental changes in *TMEM106B* expression have effects on cell toxicity (see **Figure 7**). Thus, we provide molecular characterization of the *TMEM106B* locus, elucidating a mechanism by which a causal variant at this locus may exert downstream effects on risk for FTLD.

**Figure 7:**
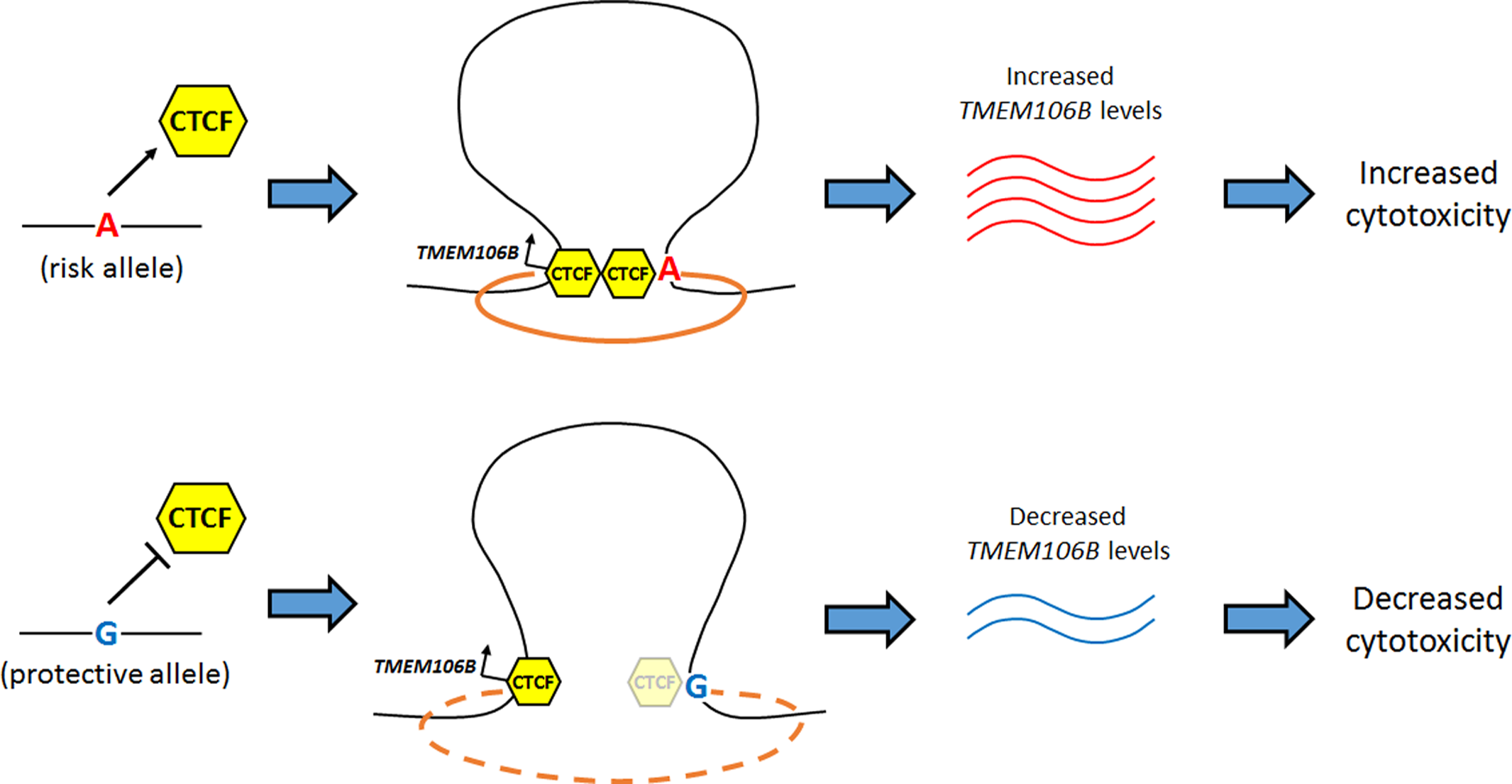
Working model of molecular mechanism underlying 7p21 association with neurodegeneration. The risk-associated allele of the causal variant (rs1990620) preferentially recruits CTCF, resulting in haplotype-specific effects on long-range chromatin interactions, with downstream effects of increased *TMEM106B* expression. Increased *TMEM106B* expression leads to increased cytotoxicity and corresponding risk for neurodegeneration.

We note that the effects we describe are incremental, rather than all-or-none. Similar incremental effects have been reported for many cases of allele-specific expression^7^,^20^,^49^,^56^,^72^^−^^74^, including allele-specific expression differences associated with neuropsychiatric disease variants^7^,^75^. Conceptually, such an incremental effect is not surprising for common genetic variants that confer only slightly increased odds for developing a disease, yet may still shed light on important disease mechanisms.

Aspects of the work presented here may be more broadly applicable to common variant effects on risk for many neurodegenerative diseases or, even more broadly, for many common variant-trait associations. For example, here, genotype at the causal noncoding variant rs1990620 alters FTLD risk through an incremental effect on *TMEM106B* expression. The enrichment of disease-associated variants in predicted *cis*-regulatory regions^42^,^43^ and the overlap between these variants and variants associated with gene expression levels (eQTLs)^43^,^76^^−^^78^ suggest that many common variants identified by GWAS may act by modulating gene expression. Consistently, most post-GWAS functional studies have identified putative causal variants that are thought to affect disease risk by altering the expression of nearby or distant genes^7^,^44^^−^^46^,^48^,^49^,^79^,^80^. However, while most of the proposed causal variants in these studies are located in regions with enhancer-associated features, our results, to our knowledge, represent the first characterization of a putative GWAS causal variant located in an architectural CTCF site. Furthermore, while GWAS causal variants have previously been implicated in specific allele-specific long-range chromatin interactions^47^,^81^^−^^84^, here we report a variant that may directly affect higher-order chromatin architecture. We also find that SNPs associated with risk for neurodegeneration by GWAS are enriched in brain CTCF binding sites, suggesting that allele-specific modulation of gene expression programs influenced by CTCF may underlie additional risk factors for other neurodegenerative diseases.

We note that the degree of molecular precision provided here is largely absent in the characterization of neurodegenerative disease loci first discovered by GWAS. Yet this level of mechanistic detail illuminating the genetic regulation and biological function of GWAS-derived loci is certainly needed if we are to translate the thousands of “leads” obtained in this way into potential avenues for therapeutic interventions. In this context, the strategy illustrated here of prioritization of variants based on the wealth of newly available genomic data, followed by targeted investigation in cell culture systems, may be more broadly applicable to the study of common variants associated with other human diseases.

## ACKNOWLEDGMENTS

We would like to thank Dr. Jonathan Schug of the University of Pennsylvania for conceptual and technical expertise in high-throughput sequencing experiments, and Nathaniel D. Berkowitz for computational assistance.

## SUPPLEMENTARY MATERIALS

**Figure S1.**
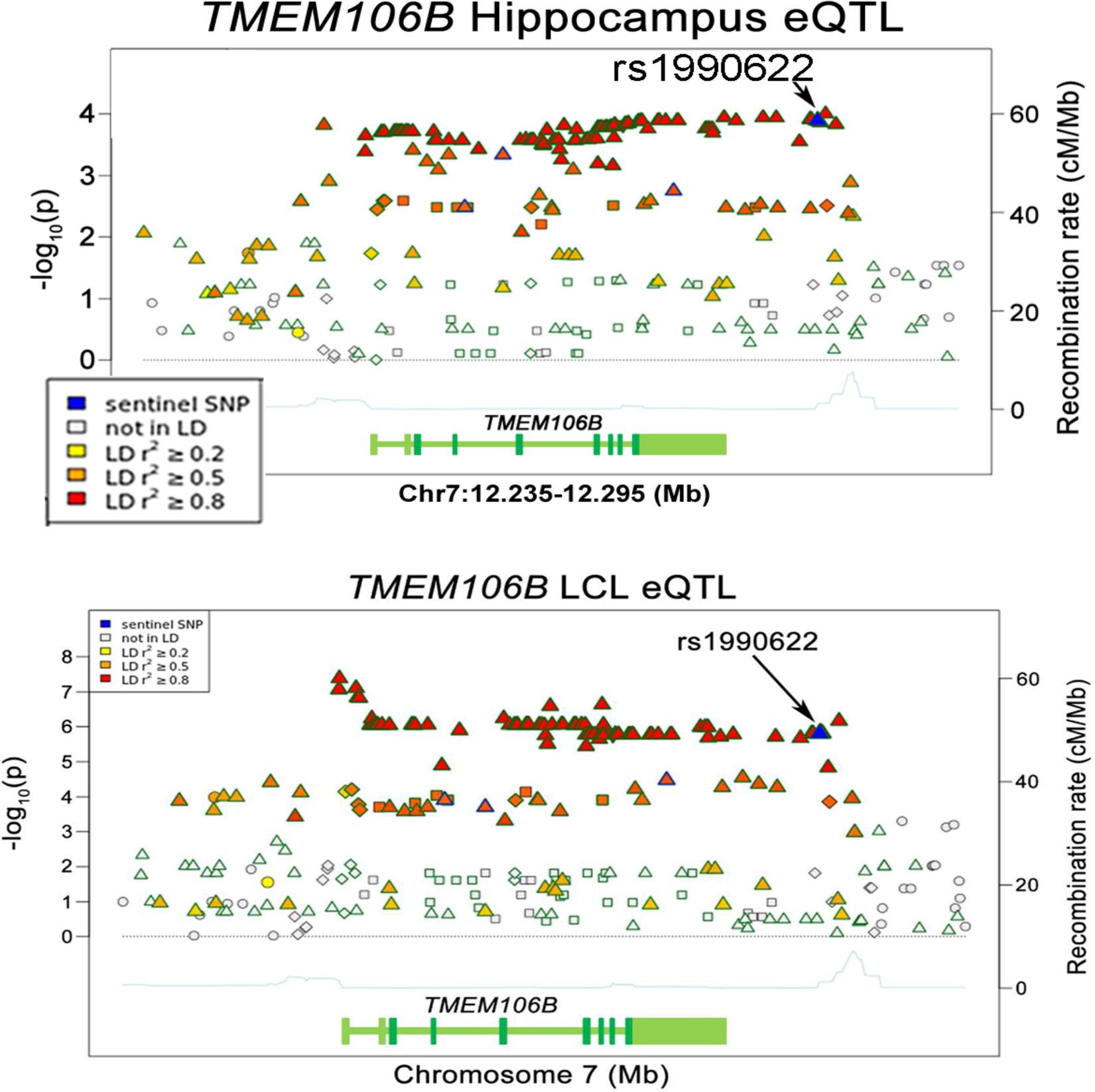
Multiple linked 7p21 variants are associated with *TMEM106B* expression. All GTEx SNPs tested for association with *TMEM106B* levels for human hippocampus (top) or LCLs (bottom) within a 60kB window are shown. SNPs are color coded according to linkage disequilibrium with rs1990622, the GWAS sentinel SNP (shaded in blue and indicated with arrow). X-axis indicates genomic position with *TMEM106B* gene structure indicated as a reference (green); y-axes indicate degree of association with *TMEM106B* levels (left) and recombination (right), indicated by blue line under eQTL plot).

**Figure S2.**
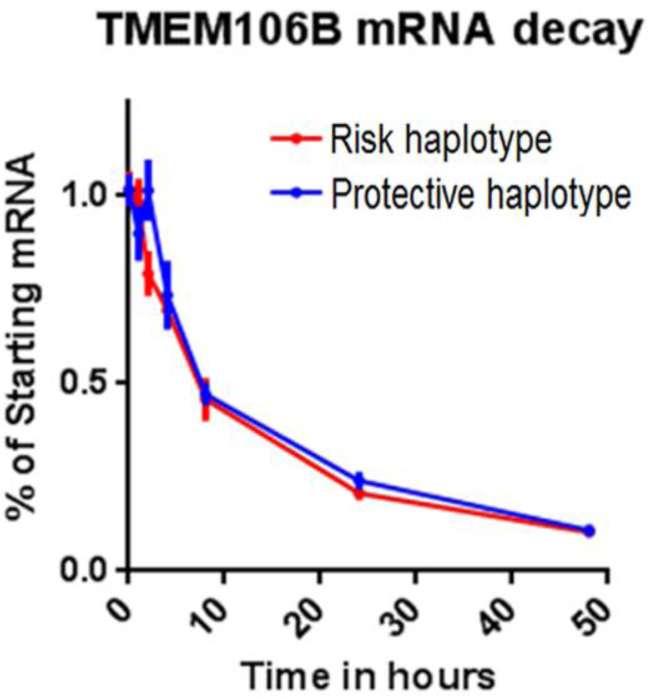
*TMEM106B* genotype does not affect mRNA stability. Lymphoblastoid cell lines homozygous for the risk (n=3) and protective (n=3) *TMEM106B* haplotypes were treated with Actinomycin D to inhibit transcription. RNA was isolated at 0h, 1 h, 2h, 4h, 8h, and 24h after treatment, and *TMEM106B* was quantified by RT-qPCR. Data were analyzed with a two-way ANOVA.

**Figure S3.**
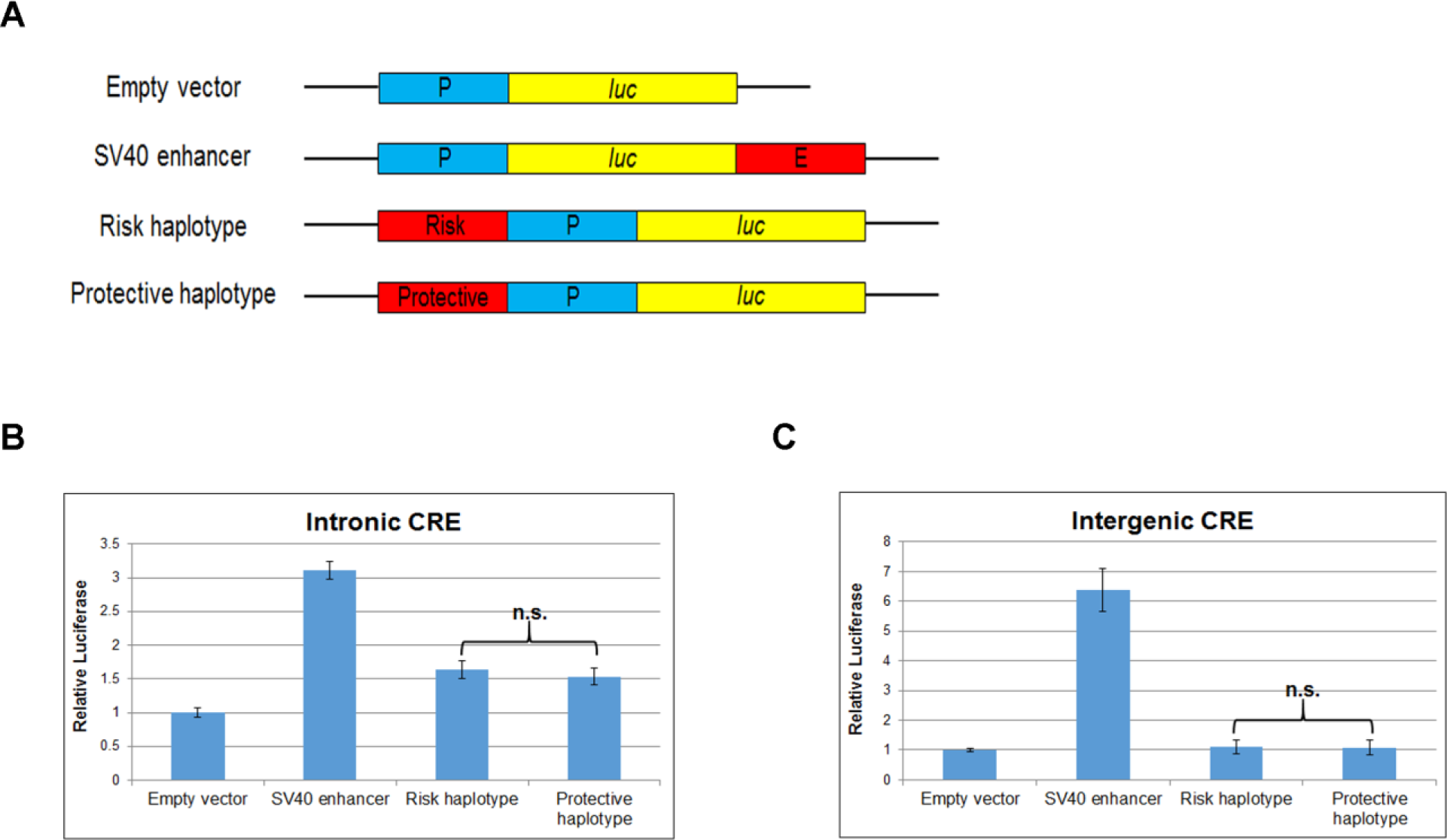
Intronic and intergenic candidate *cis*-regulatory elements (CBEs) do not harbor functional variants. (A) Schematic of luciferase reporter constructs used to test candidate CREs for enhancer activity. Empty vector construct contains no enhancer (negative control); SV40 enhancer construct serves as a positive control; test constructs contain putative CREs with either the risk or protective SNP alleles cloned upstream of the SV40 promoter. P=SV40 promoter; *luc*=luciferase; E=SV40 enhancer. (B,C) The intronic CRE displays weak ~1.5-fold enhancer activity with both SNP haplotypes (p<0.001), whereas neither SNP haplotype of the intergenic CRE displays any activity. n. s.=non-significant.

**Figure S4.**
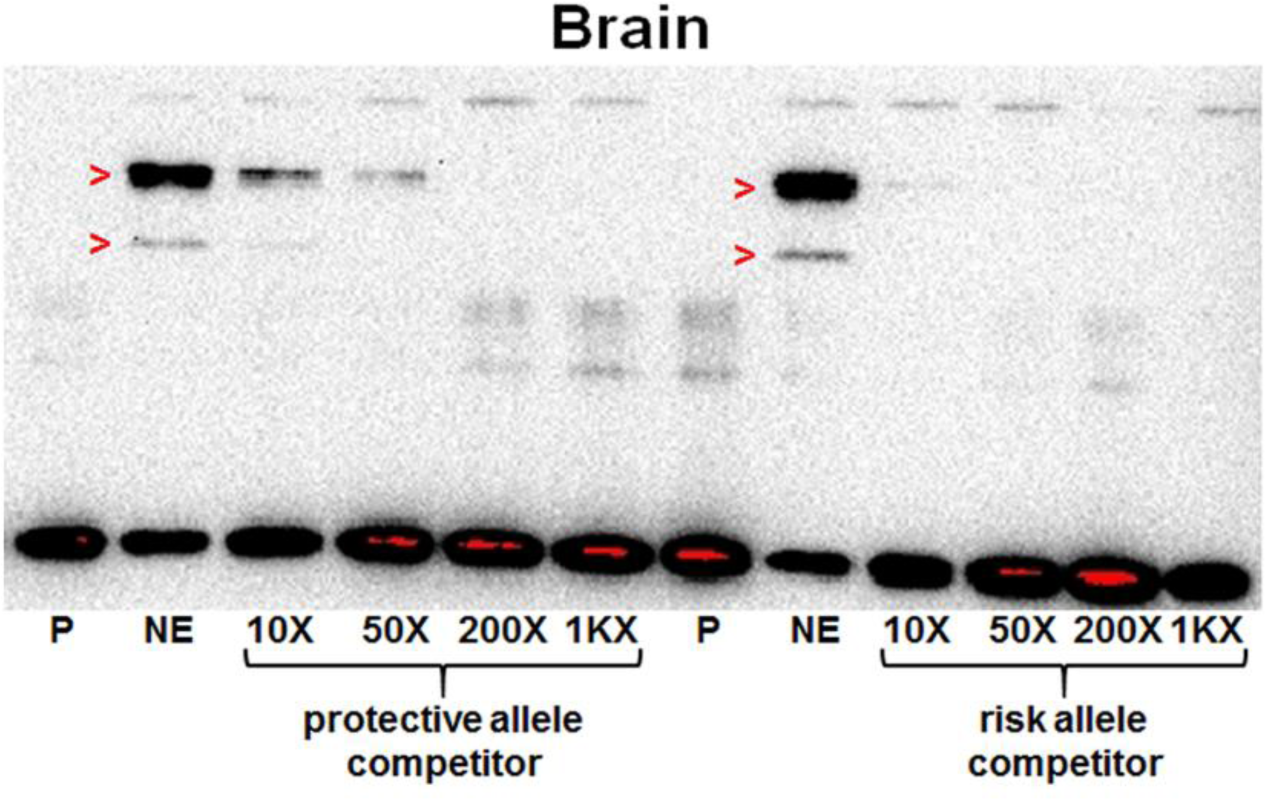
The risk allele of rs1990620 preferentially recruits a nuclear factor in brain nuclear extract. A 5’ biotinylated probe containing the rs1990620 protective allele was incubated with human brain nuclear extract, resulting in a similar shift (red arrowheads) to that seen with the rs1990620 risk allele (**Figure** **4E**). The shifted complex was better competed with excess amounts of unlabeled oligo containing the risk allele of rs1990620 than that containing the protective allele. P=probe, NE=probe+nuclear extract, 1KX=1,000X.

**Figure S5.**
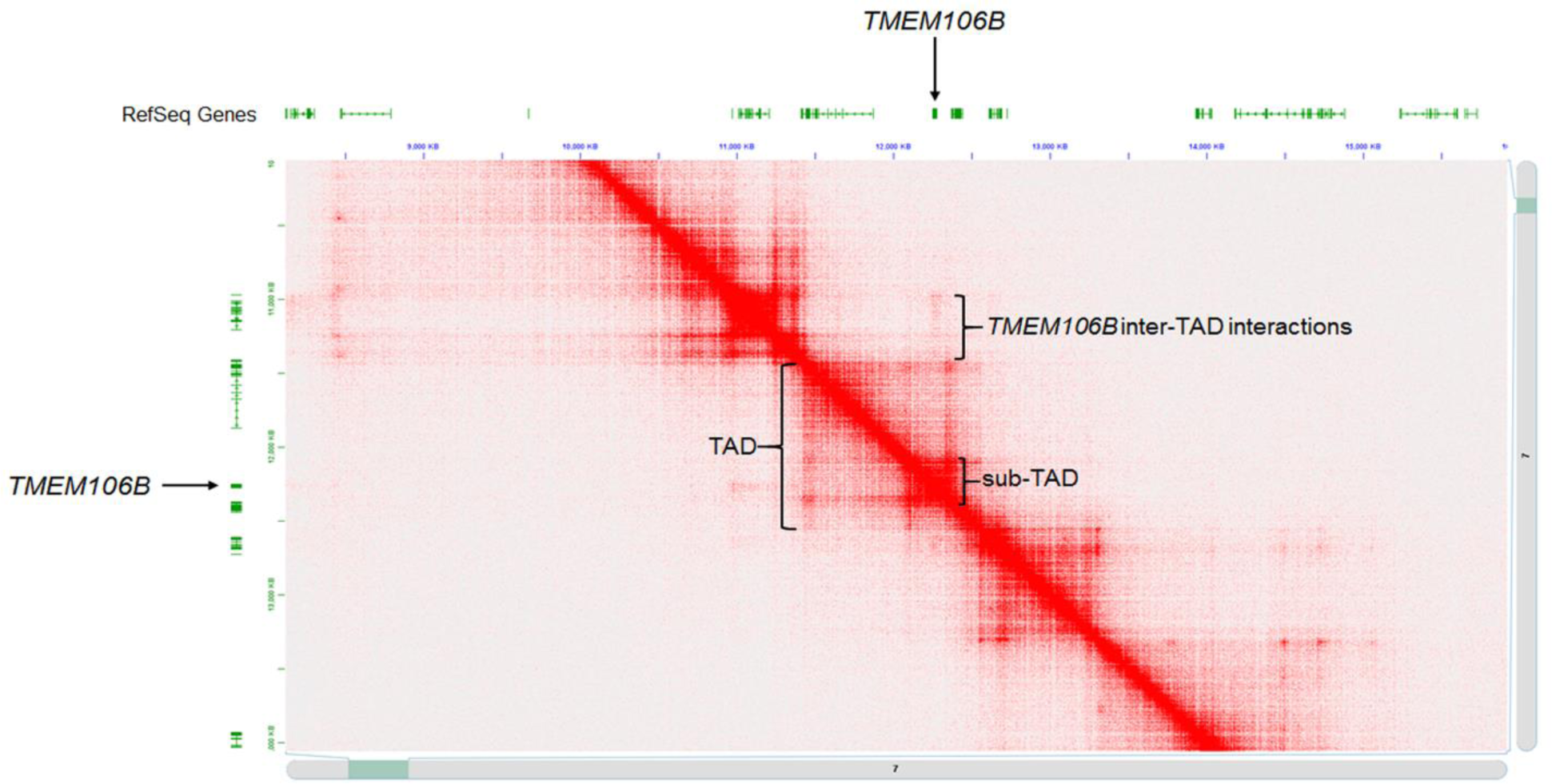
Lymphoblastoid cell line *in situ* Hi-C data reveals the chromatin architecture at the *TMEM106B* locus. Hi-C heatmap for chromosome 7p21 displays a ~250kb topologically associating domain (sub-TAD) within a larger ~1Mb TAD, containing *TMEM106B* and no other genes. *TMEM106B* also interacts with genomic regions extending several hundred kilobases upstream of the TAD (inter-TAD interactions).

**Figure S6.**
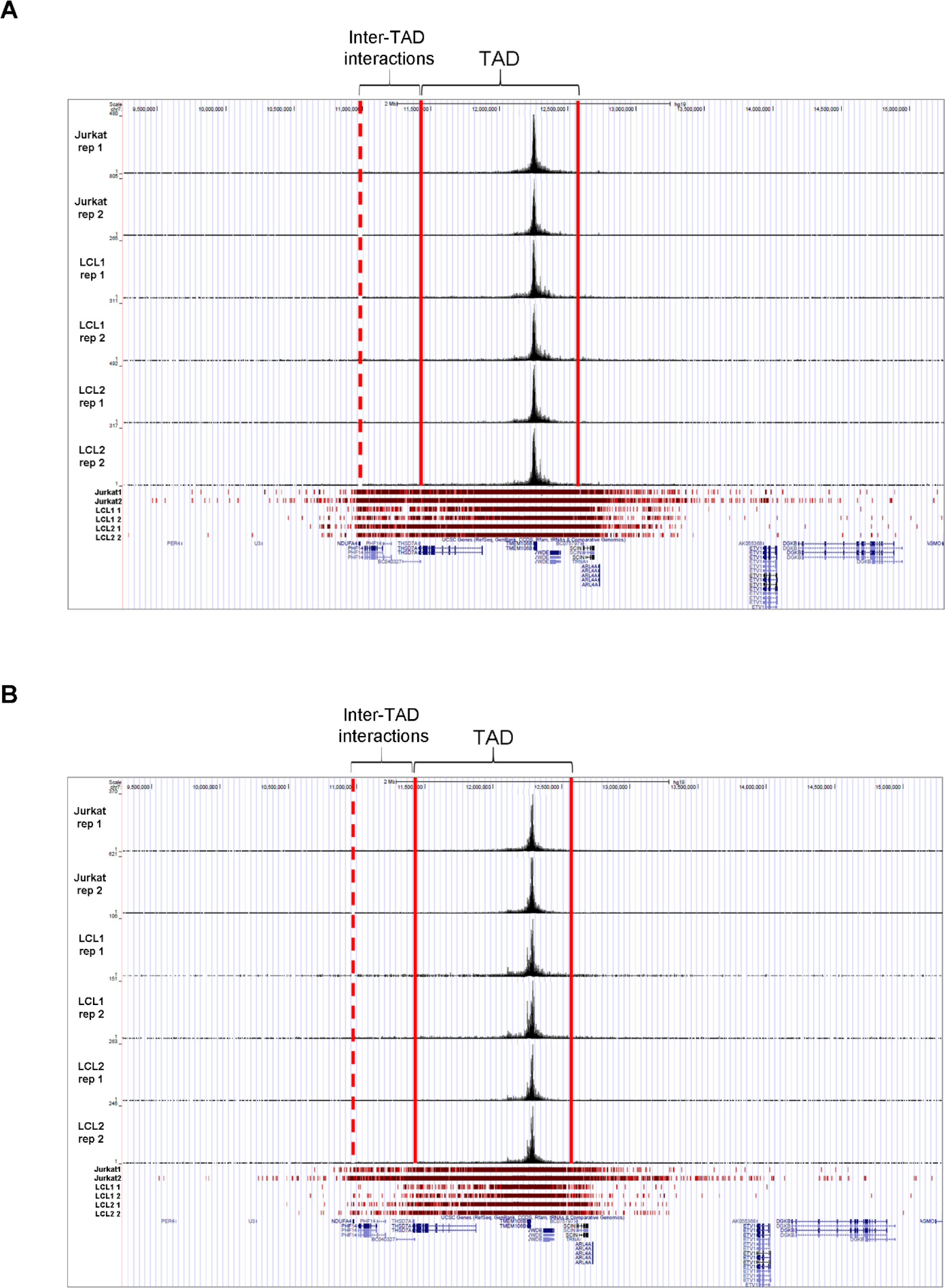
Capture-C recapitulates the predicted chromatin architecture at the *TMEM106B* locus. (A) When analyzing all long-range interactions mapping to chromosome 7 from the *TMEM106B* promoter viewpoint, statistically significant interactions are limited mostly to coordinates agreeing with the Hi-C TAD (solid red lines) and upstream inter-TAD interactions (dashed red line indicates endpoint of interactions observed in the Hi-C data). (B) Similar results are seen from the CTCF site viewpoint. The top of each panel shows raw read coverage for each sample and replicate, and the bottom of each panel shows statistically significant interactions (red bars indicate statistical significance, with darker shades of red indicating higher confidence interactions). Data are viewed in the UCSC Genome Browser, with read counts on the y-axis of each track.

**Figure S7.**
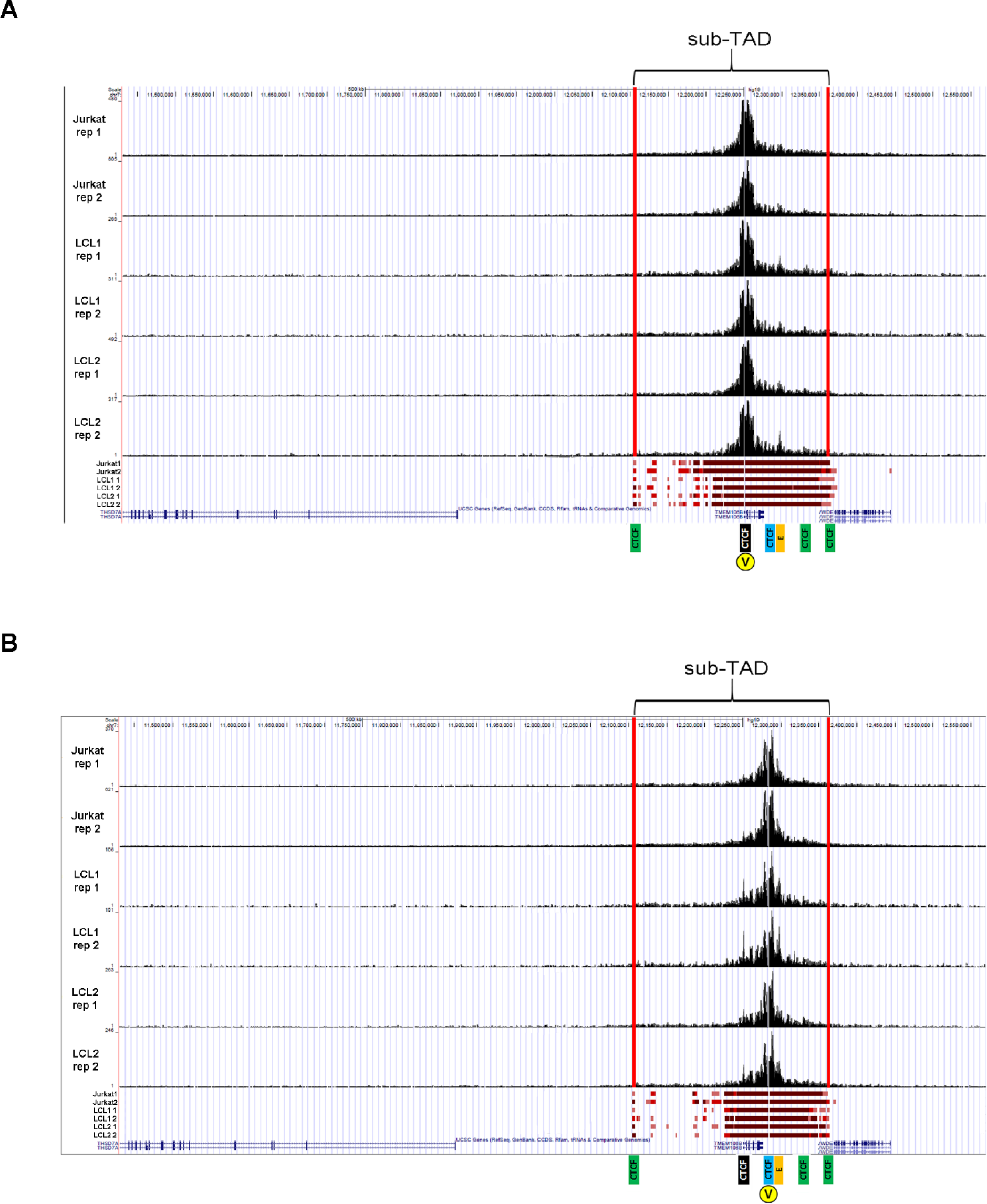
Capture-C identifies key interactions between *cis*-regulatory elements at the *TMEM106B* locus. (A) When analyzing only the interactions mapping to the TAD from the *TMEM106B* promoter viewpoint, statistically significant interactions map entirely to the Hi-C sub-TAD coordinates (solid red lines). (B) Similar results are seen from the CTCF site viewpoint. Key interacting regions from **Figure** **5A** are indicated below each browser snapshot. Yellow circles marked with a “V” indicate the Capture-C viewpoint. Data are viewed in the UCSC Genome Browser, with read counts on the y-axis of each track.

**Table S1.** Functional annotations of tested top eQTL SNPs variants. The 7 SNPs overlapping either DNase hypersensitivity (HS), transcription factor (TF) binding, or active histone marks in lymphoblastoid cell lines (LCLs) are listed by location. CTCF binding at the region containing the GWAS sentinel SNP, rs1990622, and two other linked SNPs, is seen in brain cell types as well as LCLs.

| Putative CRE | rsid | hg19 | DNase HS |  | TF binding |  | Histone marks |  | LD with rs1990622 (r<sup>2</sup>) |
| --- | --- | --- | --- | --- | --- | --- | --- | --- | --- |
|  |  |  | LCL | Brain | LCL | Brain | LCL | Brain |  |
| intronic CRE | rs1435527 | 12262571 | Yes | No | NFIC, RUNX3, NFYB | No | H3K4me1 | No | 0.96 |
| intronic CRE | rs1435525 | 12262717 | Yes | No | NFIC, RUNX3, NFYB | No | H3K4me1 | No | 0.96 |
| intronic CRE | rs1435524 | 12262801 | Yes | No | NFIC, RUNX3, NFYB | No | H3K4me1 | No | 0.96 |
| intergenic CRE | rs1548884 | 12279761 | No | No | PU.1 | No | No | No | 0.98 |
| CTCF site | rs1990622 | 12283787 | No | Yes | CTCF | CTCF | No | No | N/A |
| CTCF site | rs1990621 | 12283873 | No | Yes | CTCF | CTCF | No | No | 0.99 |
| CTCF site | rs1990620 | 12284008 | No | Yes | CTCF | CTCF | No | No | 1 |

**Table S3.** Cell lines used for CTCF ChIP-seq and DNase digital genomic footprinting analyses. All cell lines were confirmed to be heterozygous at rs1990620 by analyzing reads containing rs1990620. Major and minor allele rs1990620 reads were enumerated and pooled across samples for each experiment. Deviation from an expected ratio of 50:50 was tested using a two-tailed binomial sign test. Cell lines are listed by their ENCODE identifiers, or spelled out if no identifiers exist.

| Experiment | Cell lines |
| --- | --- |
| CTCF ChIP-seq | BE2_C, Caco-2, GM06990, GM12864, GM12872, GM12873, GM12874, GM12891, GM12892, GM19238, H1-hESC, HBMEC, HCFA, HEK293, HepG2, HRPEpiC, HUVEC, HVMF, NHDF, RPTEC |
| DNase digital genomic footprinting | AoAf, HCFaa, NHDF-neo, HMVEC-dBl-Ad, H7-hESC, HMVEC-dLy-Neo |

**Table S4.**
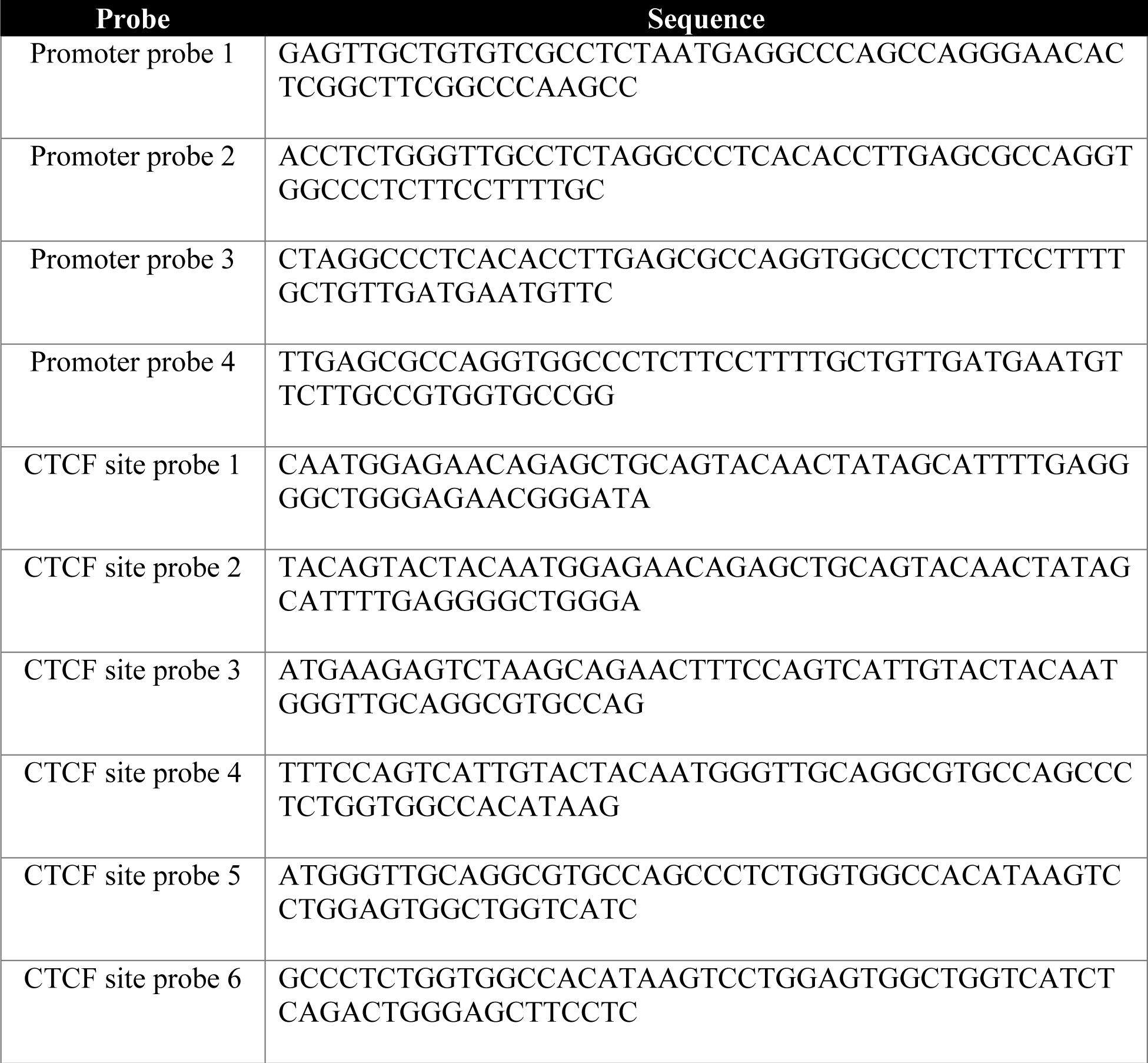
Probe sequences used for Capture-C. Four and six 60bp biotinylated probes were used to capture interactions involving the *TMEM106B* promoter and rs1990620-containing CTCF site, respectively.

